# Homology- and coevolution-consistent structural models of bacterial copper-tolerance protein CopM support a ‘metal sponge’ function and suggest regions for metal-dependent protein-protein interactions

**DOI:** 10.1101/013581

**Authors:** Luciano A. Abriata

**Affiliations:** Laboratory for Biomolecular Modeling, Swiss Federal Institute of Technology (EPFL) and Swiss Institute of Bioinformatics, 1015 Lausanne, Switzerland

**Keywords:** copper tolerance, copper resistance, copper homeostasis, CopM, molecular modeling, homology modeling, coevolution, EVFold, I-TASSER

## Abstract

Copper is essential for life but toxic, therefore all organisms control tightly its intracellular abundance. Bacteria have indeed whole operons devoted to copper resistance, with genes that code for efflux pumps, oxidases, etc. Recently, the CopM protein of the CopMRS operon was described as a novel important element for copper tolerance in *Synechocystis*. This protein consists of a domain of unknown function, and was proposed to act as a periplasmic/extracellular copper binder. This work describes a bioinformatic study of CopM including structural models based on homology modeling and on residue coevolution, to help expand on its recent biochemical characterization. The protein is predicted to be periplasmic but membrane-anchored, not secreted. Two disordered regions are predicted, both possibly involved in protein-protein interactions. The 3D models disclose a 4-helix bundle with several potential copper-binding sites, most of them largely buried inside the bundle lumen. Some of the predicted copper-binding sites involve residues from the disordered regions, suggesting they could gain structure upon copper binding and thus possibly modulate the interactions they mediate. All models are provided as PDB files in the Supporting Information and can be visualized online at http://lucianoabriata.altervista.org/modelshome.html **Note (January 2017):** Recent X-ray structures of apo, copper- and silver-bound CopM are < 3Å RMSD away from the models, and reveal metal-dependent structural flexibility (Zhao et al *Acta Crystallogr D Struct Biol.* 2016)

---

Copper is an element essential for life, but it becomes toxic at concentrations exceeding organismal requirements.^1,2^ Therefore, living creatures have evolved complex homeostatic systems that monitor copper concentrations and manage its intake and distribution to the proteins that require it, and that prevent its toxic effects.^1,3^ Bacteria carry whole genetic operons and regulons devoted to conferring tolerance against Cu(I) and Cu(II). The proteins coded in these genetic elements achieve protective roles through varied mechanisms, including efflux systems, reduction of the more toxic Cu(I) to Cu(II), metal chelation and other less understood functions.^4–9^

Recently, a novel protein dubbed CopM was identified in the *copMRS* operon of a copper-resistance regulon of *Synechocystis*.^10,11^ The CopM protein is of periplasmic and extracelullar localizations,^11^ and plays an important role in resistance according to the effects observed in knock out strains.^11^ *In vitro,* it can tightly bind one equivalent of Cu(I) or, less tightly, of Cu(II).^11^ However, its amino acid sequence includes several histidine and methionine residues that could allow it to bind more metal equivalents.^12^ Indeed, other periplasmic copper-tolerance proteins that function as metal sponges, like *E. coli* PcoE, retain one metal ion upon purification but accept more on titration.^13^

This work reports structural models of CopM based on classical homology modeling and on a protocol that uses evolutionary couplings to model a protein’s 3D structure. These two methods, based on fundamentally different concepts and data, lead to essentially the same structural features. Specifically, CopM is predicted a 4-helix bundle that brings together several His and Met side chains into proximity, giving place to many potential copper-binding sites, with two disordered regions that could become ordered upon copper binding. Together with basic bioinformatic analyses of CopM’s sequence, the emerging model supports the hypothesis of the protein working as a copper sponge, possibly to rapidly buffer sudden increases in environmental copper, with the potential additional function of differentially interacting with other proteins depending on its metallation level, possibly to modulate responses beyond initial chelation.

## Results and Discussion

*Analysis of Synechocystis CopM sequence.* The amino acid sequence of full CopM from *Synechocystis* sp. PCC6803 (Uniprot ID Q55943_SYNY3) is shown in Fig. 1. It consists of a DUF305 domain (domain-of-unknown function number 305, PF03713, underlined in Fig. 1). The sequence begins with a short hydrophobic segment (italics in Fig. 1) previously regarded as a signal peptide for excretion; however, a computational prediction specifically designed to tell transmembrane regions from signal peptides suggests that this is a transmembrane helical region.^14^ As such, the protein is predicted to be mostly in the periplasm anchored to the membrane, consistently with < 30% leaking out in the first 24 h of growth as reported.^11^

Secondary structure predictions (red in Fig. 1) disclose four main helical elements with high confidence plus two short regions of helical propensity. Disorder predictions with Disopred3^15^ (lowercase in Fig. 1) suggest two main unstructured regions, to which the short segments of helical propensity map. Both disordered regions are predicted by Disopred3 to be involved in interactions with other proteins.

**Figure 1.**
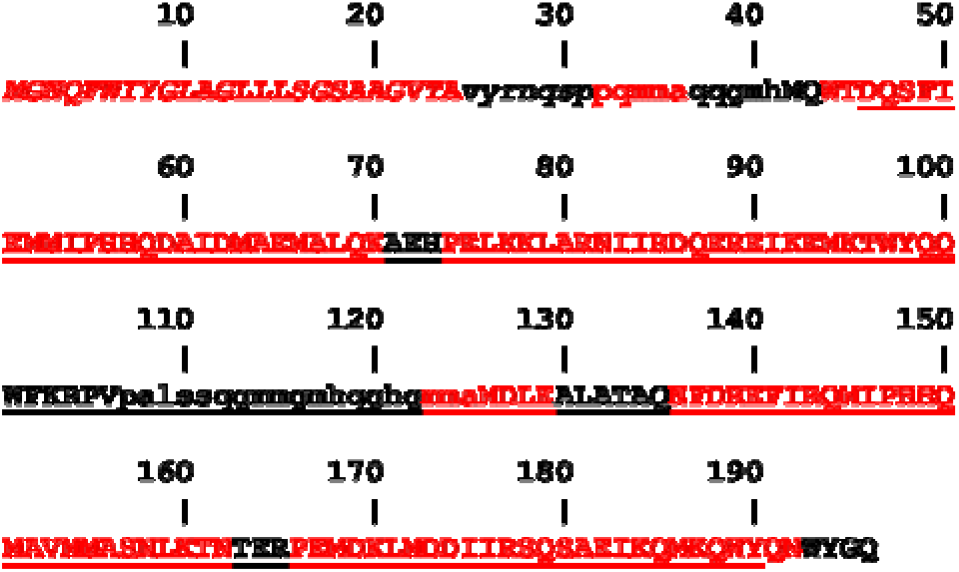
Amino acid sequence of *Synechocystis* CopM, on which bioinformatic predictions were mapped: secondary structure consensus from several programs (red=helix), SignalP’s prediction of a transmembrane helix rather than a signal peptide (italics), and disordered regions predicted by Disopred3 (lowercase).

*Homology and coevolution-based modeling.* Models of proteins and their dynamics often lead to new hypotheses and explanations. In a first exploratory step, two models of the globular domain of CopM (residues 25-196) were built using either classical homology modeling (with I-TASSER^17^) or a tool that analyzes evolutionary couplings throughout a protein’s sequence to fold it into a consistent 3D structure (EVFold^18–20^). These two protocols are completely unrelated and rely on fundamentally different data and methods. They independently produced similar models (Fig. 2 top), both of high confidence, indicating that the global features of the models should not differ largely from those of the “true” structure. The high confidence of the homology model stems from the availability in the Protein Data Bank of X-ray structures for two DUF305 domains with sequence identity > 20% to CopM (PDB ID 3BT5 and 2QF9). In turn, the high confidence of the coevolution-based prediction stems from the large number of sequences compiled by EVFold in an alignment with standard parameters (2586) that reaches good coverage of the sequence and of the amino acid space at each position of the sequence.

The largest differences between the EVFold and I-TASSER models map to the two predicted disordered regions, which are heterogeneous themselves even among the different models produced by each program. This is likely due to a combination of true dynamics, lack of homolog elements in the X-ray structures used by I-TASSER, and a low sequence complexity that prevents extraction of direct couplings by EVFold. To improve the definition of these regions, a new set of models was built by feeding the coevolution information from EVFold into I-TASSER. This procedure results in four similar top models (pairwise RMSDs between 0.9 and 4.1 Å, Z-scores from -1.61 to -0.51) that are better packed than the EVFold-only model and that reproduce the EVFold contacts better than the I-TASSER-only model. These models show variability mainly in the first 15-20 N-terminal residues (Fig. 2 bottom and Fig. 3).

**Figure 2.**
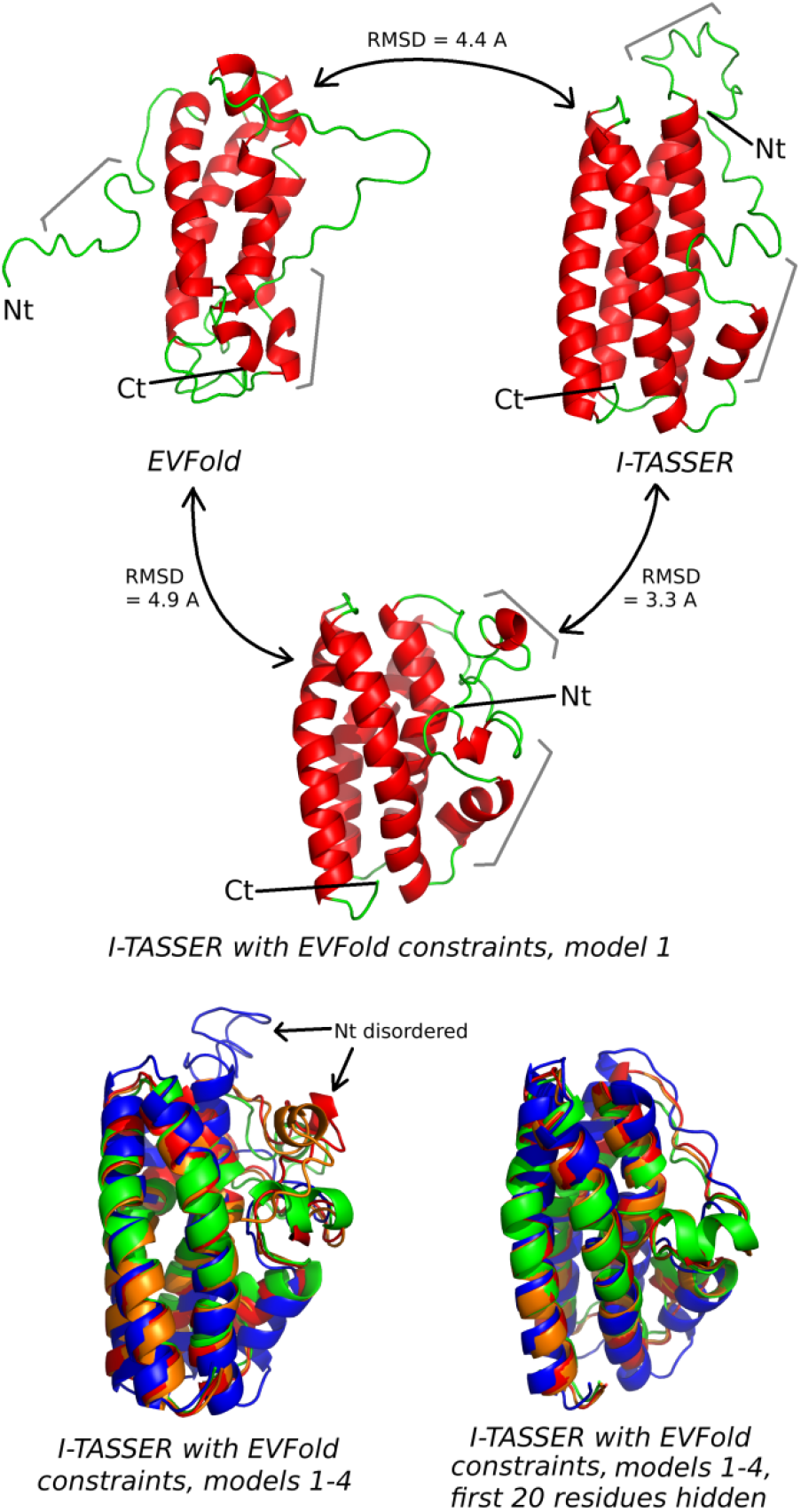
Models of CopM produced by EVFold (top left), I-TASSER (top right), and I-TASSER supplemented with EVFold-derived restraints (middle, first model only; and bottom, four best models, full on the left and without the first 20 residues on the right, all displayed individually in Fig. 3). All shown structures are aligned in space. Gray brackets point at the two predicted disordered regions. Black arrows in the bottom left point at the two main conformations modeled for the N-terminal region. Figures rendered with PyMOL.^16^

**Figure 3.**
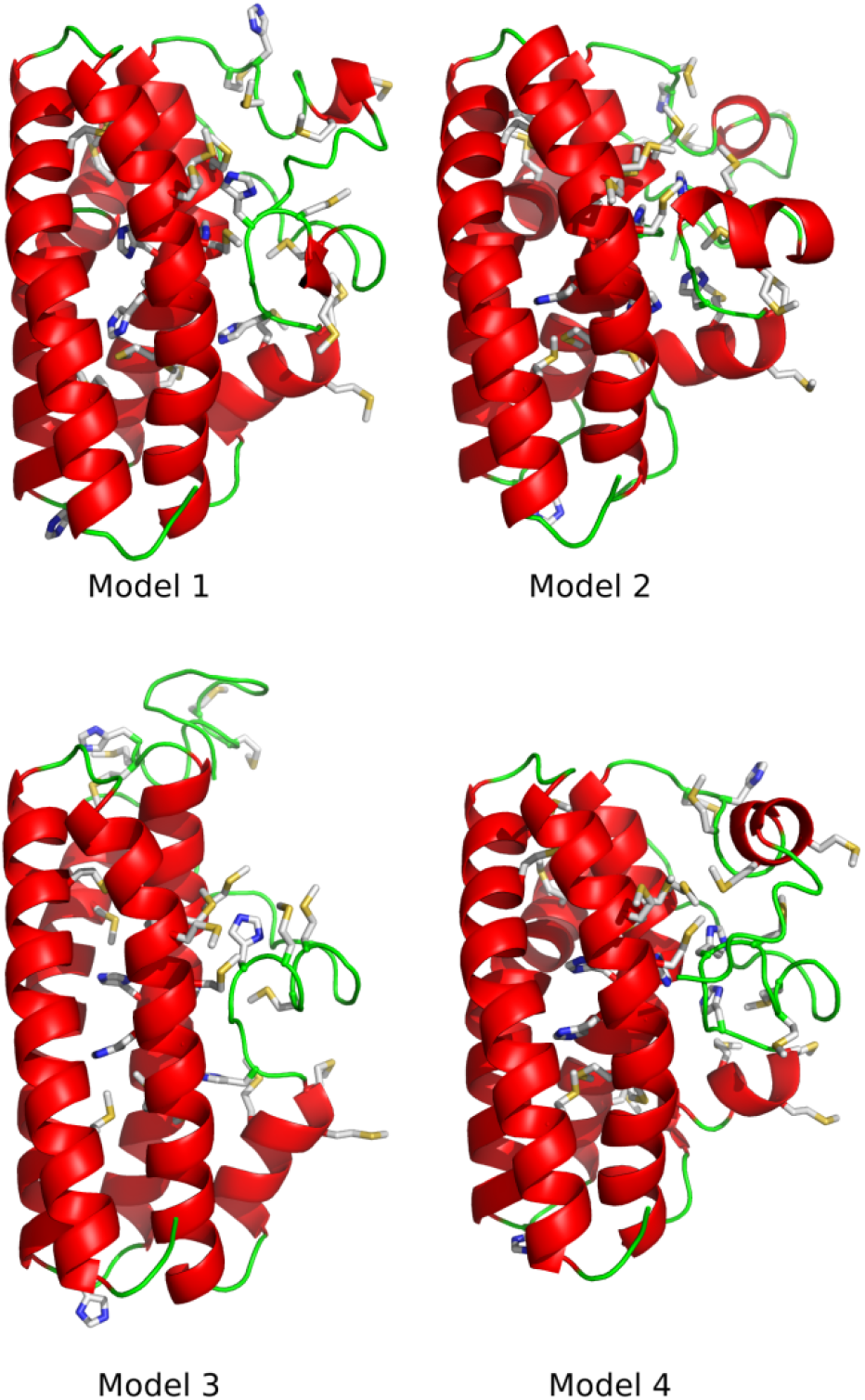
The four top models produced by combining contacts computed by EVFold with homology modeling by I-TASSER. Histidine and methionine residues are rendered as sticks with standard color codes for heavy atoms.

All the obtained models display a 4-helix bundle topology, with an overall fold that brings most Met and His side chains inside the lumen of the bundle (Fig. 3 and Fig. 4). This results in many potential metal-binding sites inside the protein, some of them rather isolated but others making a continuous network of potential sites, most remarkably a central cluster of buried histidines (Fig. 4). Also the two disordered regions form arrangements reminiscent of copper-binding sites, by combining their methionine and histidine residues with those of the 4-helix bundle.

**Figure 4.**
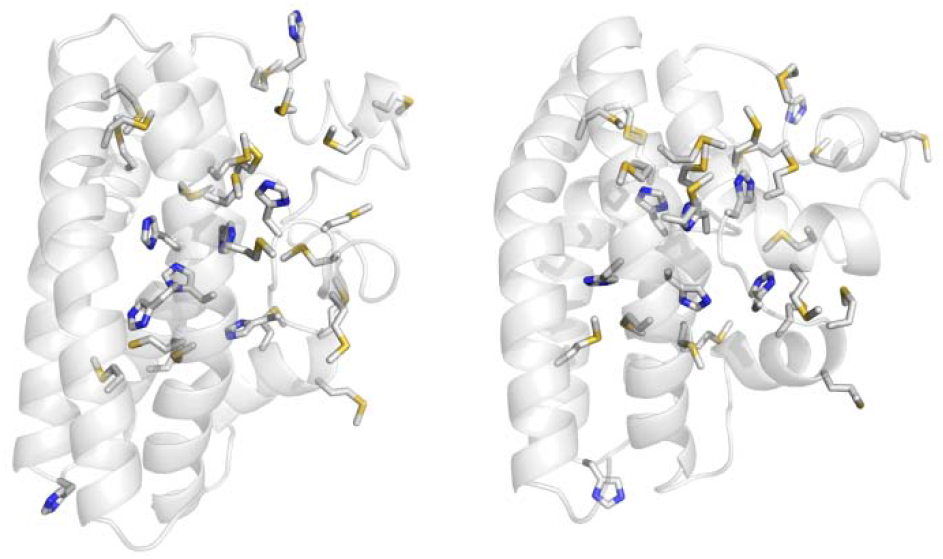
Models 1 (left) and 2 (right) from Fig. 3, with transparent cartoons to better evidence the distribution of histidine and methionine residues.

Based on the structural models and sequence predictions, it is possible that the two predicted disordered regions are truly unstructured and dynamic in the metal-free protein, and that they become folded upon copper binding, as observed for other copper-binding proteins.^21,22^ An open, flexible state in the absence of copper could facilitate metal uptake by the many internal sites of the protein. After such initial role as a “copper sponge” capable of buffering sudden increases in environmental copper, metal binding-induced structuring of the disordered regions could possibly trigger a second response by altering interactions with other proteins.

One last interesting point stems from two pairs of sequence segments that are significantly predicted to interact according to EVFold’s residue-residue couplings, but that none of the three modeling procedures could satisfy, not even the one based exclusively on EVFold data (Fig. 5). In the few examples where such situation was reported in the literature, the couplings turned out to correspond to either homodimerization surfaces^19^ or to alternative conformations such as intermediate and hidden states.^23^ Given the localization of these regions in the models, it is in this case more likely that these sequence segments mediate dimerization. An alternative conformation is less likely because it would entail very large rearrangements, although it cannot be discarded so both possibilities deserve exploration.

**Figure 5.**
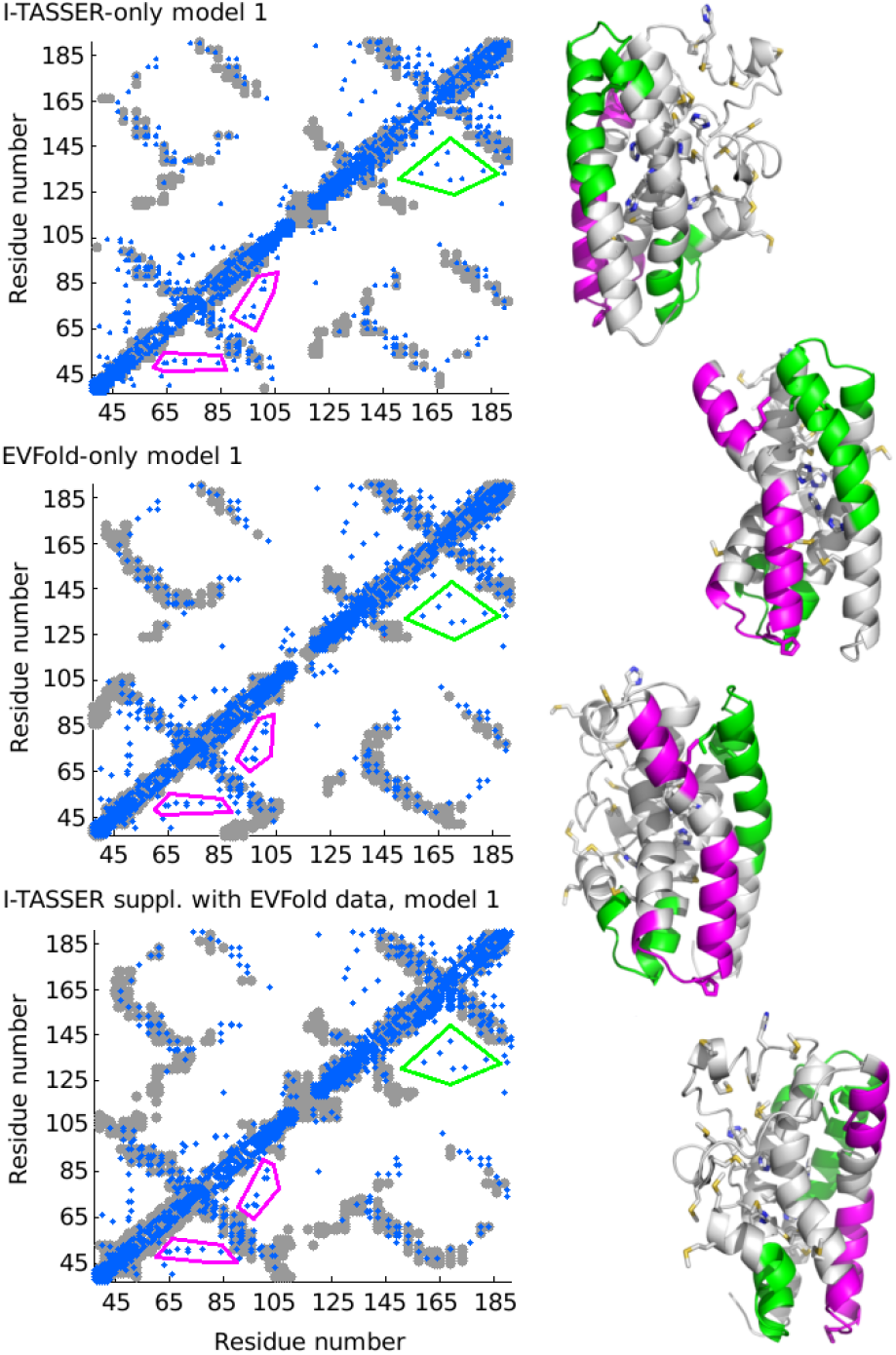
Left: overlay of EVFold-predicted residue-residue contacts (blue, PLM coupling > 0.13) on the contact maps of the best model produced by each modeling protocol. Right: Four views of model 1 produced by I-TASSER supplemented with EVFold restraints, with histidine and methionine residues rendered as sticks. Cartoon segments in magenta and green point at the two regions where the contacts predicted by EVFold are not satisfied in the models, as shown by corresponding traces in the plots on the left.

## Conclusions

Coevolution-based methods for modeling protein structures and their complexes are rapidly getting established ^18–20,24–27^ and starting to be used for real-world applications.^28,29^ Together with increasingly confident tools for homology modeling and sequence-based predictions, coevolution-based methods have an enormous potential to explain existing results and drive the generation of new hypotheses. Hopefully, the models and analyses described here will fulfill this goal, in this case headed towards a better understanding of how CopM and possibly other similar copper-tolerance proteins work.

## Supporting Information

PDB files for the six mentioned models (best model from I-TASSER, best model from EVFold, plus four models produced by I-TASSER with EVFold restraints) are provided in the Supporting Information. A short Methods section describing how the models were generated is also given.

## References

(1) Bertini, I.; Cavallaro, G.; McGreevy, K. S. Coord Chem Rev 2010, 254, 506–524.

(2) Robinson, N. J.; Winge, D. R. Annu. Rev. Biochem. 2010, 79, 537–562.

(3) Waldron, K. J.; Robinson, N. J. Nat. Rev. Microbiol. 2009, 7, 25–35.

(4) Grass, G.; Rensing, C. Biochem. Biophys. Res. Commun. 2001, 286, 902–908.

(5) Pontel, L. B.; Soncini, F. C. Mol. Microbiol. 2009, 73, 212–225.

(6) Depuydt, M.; Leonard, S. E.; Vertommen, D.; Denoncin, K.; Morsomme, P.; Wahni, K.; Messens, J.; Carroll, K. S.; Collet, J.-F. Science 2009, 326, 1109–1111.

(7) Rensing, C.; Fan, B.; Sharma, R.; Mitra, B.; Rosen, B. P. Proc. Natl. Acad. Sci. U. S. A. 2000, 97, 652–656.

(8) Abriata, L. A.; Pontel, L. B.; Vila, A. J.; Dal Peraro, M.; Soncini, F. C. J Inorg Biochem 2014, 140, 199–201.

(9) Espariz, M.; Checa, S. K.; Audero, M. E. P.; Pontel, L. B.; Soncini, F. C. Microbiol. Read. Engl. 2007, 153, 2989–2997.

(10) Giner-Lamia, J.; López-Maury, L.; Reyes, J. C.; Florencio, F. J. Plant Physiol. 2012, 159, 1806–1818.

(11) Giner-Lamia, J.; López-Maury, L.; Florencio, F. J. MicrobiologyOpen 2014.

(12) Abriata, L. A. Acta Crystallogr. D Biol. Crystallogr. 2012, 68, 1223–1231.

(13) Zimmermann, M.; Udagedara, S. R.; Sze, C. M.; Ryan, T. M.; Howlett, G. J.; Xiao, Z.; Wedd, A. G. J. Inorg. Biochem. 2012, 115, 186–197.

(14) Petersen, T. N.; Brunak, S.; von Heijne, G.; Nielsen, H. Nat. Methods 2011, 8, 785–786.

(15) Jones, D. T.; Cozzetto, D. Bioinforma. Oxf. Engl. 2014.

(16) DeLano, W. L. The PyMOL Molecular Graphics System; DeLano Scientific, San Carlos, CA, 2002.

(17) Roy, A.; Kucukural, A.; Zhang, Y. Nat. Protoc. 2010, 5, 725–738.

(18) Marks, D. S.; Hopf, T. A.; Sander, C. Nat. Biotechnol. 2012, 30, 1072–1080.

(19) Hopf, T. A.; Colwell, L. J.; Sheridan, R.; Rost, B.; Sander, C.; Marks, D. S. Cell 2012, 149, 1607–1621.

(20) Marks, D. S.; Colwell, L. J.; Sheridan, R.; Hopf, T. A.; Pagnani, A.; Zecchina, R.; Sander, C. PloS One 2011, 6, e28766.

(21) Zaballa, M.-E.; Abriata, L. A.; Donaire, A.; Vila, A. J. Proc. Natl. Acad. Sci. U. S. A. 2012, 109, 9254–9259.

(22) Abriata, L. A.; Vila, A. J.; Dal Peraro, M. J Biol Inorg Chem 2014, 19, 565–575.

(23) Morcos, F.; Jana, B.; Hwa, T.; Onuchic, J. N. Proc. Natl. Acad. Sci. U. S. A. 2013, 110, 20533–20538.

(24) Sulkowska, J. I.; Morcos, F.; Weigt, M.; Hwa, T.; Onuchic, J. N. Proc. Natl. Acad. Sci. U. S. A. 2012, 109, 10340–10345.

(25) Kamisetty, H.; Ovchinnikov, S.; Baker, D. Proc. Natl. Acad. Sci. U. S. A. 2013, 110, 15674–15679.

(26) Ovchinnikov, S.; Kamisetty, H.; Baker, D. eLife 2014, 3, e02030.

(27) Hopf, T. A.; Schärfe, C. P. I.; Rodrigues, J. P. G. L. M.; Green, A. G.; Kohlbacher, O.; Sander, C.; Bonvin, A. M. J. J.; Marks, D. S. eLife 2014, 3.

(28) Tian, P.; Boomsma, W.; Wang, Y.; Otzen, D. E.; Jensen, M. H.; Lindorff-Larsen, K. J. Am. Chem. Soc. 2015, 137, 22–25.

(29) Wickles, S.; Singharoy, A.; Andreani, J.; Seemayer, S.; Bischoff, L.; Berninghausen, O.; Soeding, J.; Schulten, K.; van der Sluis, E. O.; Beckmann, R. eLife 2014, 3, e03035.

